# Listener’s vmPFC simulates speaker choices when reading between the lines

**DOI:** 10.1101/800359

**Authors:** Qingtian Mi, Cong Wang, Colin F. Camerer, Lusha Zhu

## Abstract

Humans possess a remarkable ability to understand what is and is not being said by conservational partners. An important class of models hypothesize that listeners decode the intended meaning of an utterance by assuming speakers speak cooperatively, simulating the speaker’s rational choice process and inverting this process for recovering the speaker’s most probable meaning. We investigated whether and how rational simulations of speakers are represented in the listener’s brain, when subjects participated in a referential communication game inside fMRI. In three experiments, we show that listener’s ventromedial prefrontal cortex encodes the probabilistic inference of what a cooperative speaker should say given a communicative goal and context. The listener’s striatum responds to the amount of update on the intended meaning, consistent with inverting a simulated mental model. These findings suggest a neural generative mechanism subserved by the frontal-striatal circuits that underlies our ability to understand communicative and, more generally, social actions.

## Main Text

A cornerstone of effective communication is our ability to read between the lines, or to recognize the intended meaning of a speaker, even when the meaning is not coded in the utterance directly. The process of disambiguating the intended meaning of a speaker, often known as pragmatic interpretation, has long been hypothesized to rely on cooperation between communicators: A speaker tailors an utterance to help a listener recognize a meaning, and a listener recovers the intended meaning by assuming that the speaker spoke to be understood (*1*). The hypothesized role of cooperation has inspired a wealth of philosophical inquiries (*2*-*5*) and, more recently, empirical (*6*-*8*) and computational (*9, 10*) investigations into human communication, but little is known about the link between the key computational principles and underlying neural mechanisms.

Computationally, pragmatic interpretation requires a listener to identify the speaker’s underlying intention that motivated the choice of the utterance. According to an important class of models, this can be achieved through an internal generative process similar to how the brain translates sensations into perceptions (*10*-*13*). Decades of work suggests that the brain infers sensory causes (e.g., an object) from their bodily effects (e.g., a retinal image) by modeling the sensation-generating process and then inverting this model to derive the most probable cause of the sensation (*11, 14, 15*). In pragmatic interpretation, a similar strategy for a listener entails modeling the speaker’s decision-making process, that is, determining how an interacting web of causes–– intention, context, and knowledge––give rise to the choice of an utterance. It has been proposed that a listener simulates speaker behavior using a rational, goal-directed choice model (*10*). Speakers are expected to compare candidate expressions to make a choice for best helping the audience recognize the intended meaning in a given context. In addition, listeners need to monitor knowledge and beliefs shared with the speaker (common ground) for simulating speaker behavior based on mutual, rather than the listener’s own private knowledge (*4, 6*).

This generative account, also known as the Rational Speech Act (RSA) model (*10*), provides precise and falsifiable behavioral predictions in a parsimonious framework for social signal interpretation that can be extended from communication (*10, 12, 13*) to social perception (*16, 17*) and interpersonal decision-making (*18*). However, no direct evidence is available suggesting that pragmatic interpretation indeed involves rational, context-specific simulation of speakers. In particular, it is unclear whether the putative mental simulation signals are represented in the listener’s brain, and flexibly facilitate the utterance interpretation in varying contexts. It also remains to be explored whether the listener’s brain actively interrogates received utterances using automatically generated simulations of speakers or, given that such mental modeling is cognitively costly, produces internal estimation only when necessary (e.g., in the face of communicative ambiguity). A third unresolved question concerns how the mental simulation is internally represented––for example, whether different aspects of information (e.g., utterance, context, common ground information) directly support or modulate the putative simulation signal in the listener’s brain, or whether some aspects of information are abstracted away during the simulation process.

We addressed these questions by investigating a simple yet well-characterized game of referential communication (*7, 19, 20*), in three experimental conditions (Fig. 1A), using model-based functional magnetic resonance imaging (fMRI) (*21*). This method allowed us to connect the trial-wise brain activity during utterance interpretation with latent signals derived from computational modeling of listener behavior. A total of 41 subjects were scanned while they participated as “listeners” against a pool of anonymous “speakers”. As a first test of our hypotheses, we conducted the experiment where a listener and a speaker were randomly matched in pairs in each trial and faced the same communicative context consisting of three objects with varying colors and shapes (*symmetric condition*) (Fig. 1B and Figs. S1-2). The speaker was asked to refer to a target object by choosing between alternative expressions denoting either the color or shape of the target. The listener, who did not know the target, needed to recover the intended referent from the referring expression chosen by the speaker (*22*). Although speakers did not describe the targets in complete detail, the rate of listeners recovering the targets was as high as 75.72 ± 0.61% (mean ± SEM), significantly greater than that resulting from literal interpretations (literal recovery rate = 66.88%; *t*_40_ = 14.53, *P* < 2 × 10^−16^) (Fig. S5).

**Fig. 1.**
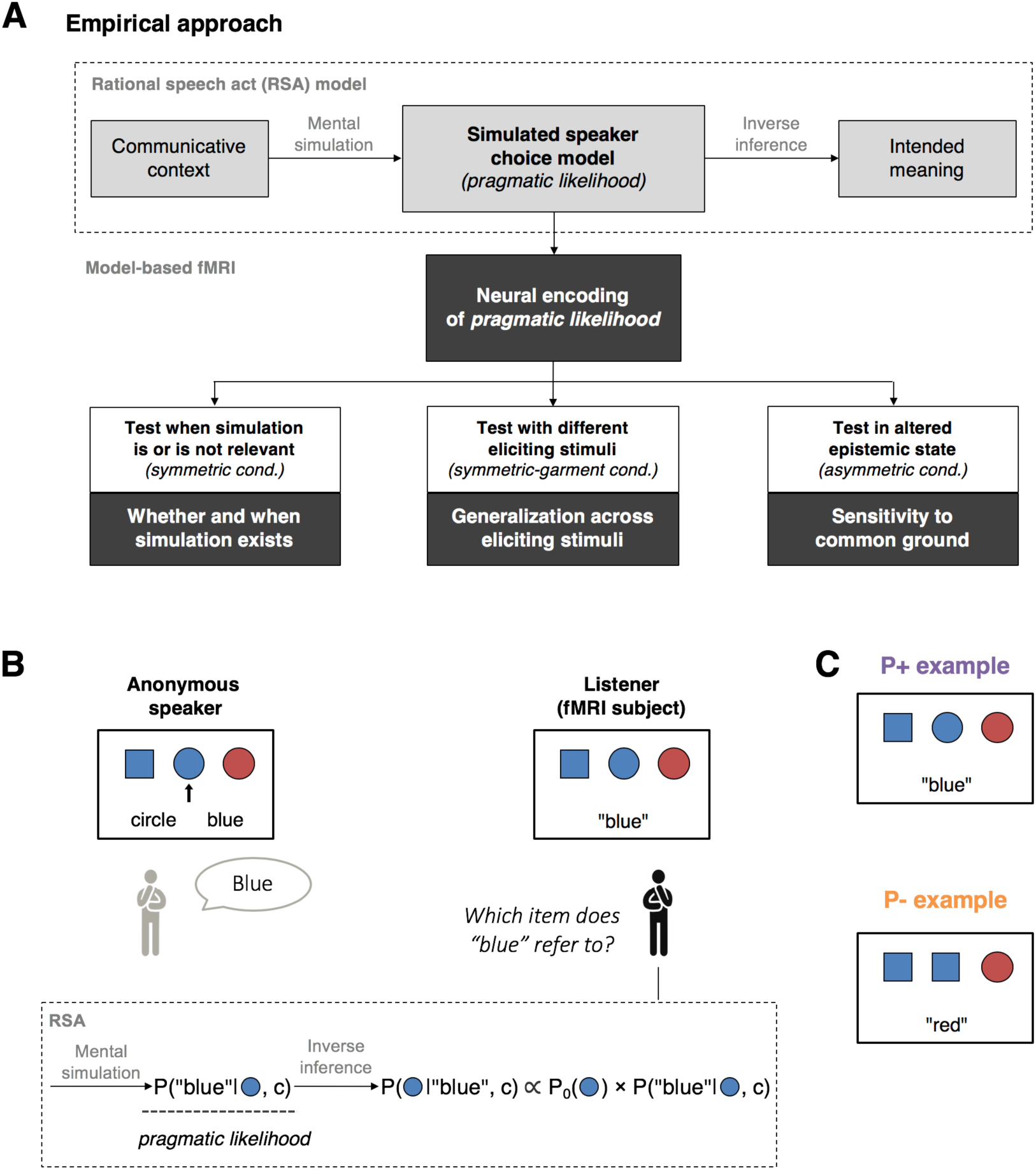
Empirical approach and task schematic. **(A)** Using model-based fMRI, we investigated the core assumption of the RSA framework, that a listener rationally simulates the speaker’s decision-making process during pragmatic interpretation. We tested this hypothesis in three communicative setups, designed for exploring the existence, generality, and sensitivity of the rational simulation signal in the listener’s brain. **(B)** Schematic of the referential game in the first experimental setup, *symmetric condition* (top), where the presence of three geometric objects (*context*) is common knowledge shared between communicators (*22*). The context and target location (indicated by the arrow in the speaker screen) varied across trials. Referential interpretation is modeled as a Bayesian inferential process using RSA (bottom): A listener evaluates how likely a particular object is to be the target given the received expression and context by inverting a mental model of speaker behavior characterized by *pragmatic likelihood* (*22*). **(C)** Trial types in the symmetric condition, differing according to whether predicting speaker choices is relevant (P+) or is not relevant (P-) for resolving communicative ambiguity. In the P+ example (top), if the target were the blue square, a speaker would have uttered “square”, which denotes the blue square unambiguously. The fact that the speaker sent “blue” instead of “square” indicates to the listener that the blue circle is the target. In contrast, in the P-example (bottom), a listener may single out the red circle upon receiving the expression “red”, without realizing that the red circle would be referred to as “red” with 50% probability by the speaker.

The paradigm has three key advantages for providing a quantitative framework connecting listener choices with the underlying cognitive processes in the context of model-based fMRI: (i) a large number of trials for each listener, (ii) tight control of the decision space of communicators, and (iii) parametric manipulation of contexts. We fitted the listener choices with the RSA model, in which listeners anticipate speakers to choose the most specific (informative) expression to refer to a target (e.g., “square” is more specific than “blue” for denoting the blue square in Fig. 1B). By comparing the specificity between competing references, listeners simulate the probability that a speaker chooses a particular reference given a target and context (henceforth *pragmatic likelihood*), and then invert pragmatic likelihood with Bayes’ rule for deriving the most probable referent (Fig. 1B, bottom) (*22*).

This model, together with the task design in which we systematically varied reference specificity through manipulating features of the presented objects (*22*), allowed us to create the trial-wise regressor of pragmatic likelihood and explore its neural encoding in the listener’s brain. Furthermore, to test the hypothesis that pragmatic likelihood is automatically registered in the listener’s brain, even when mental simulation of the speaker is not required, we classified choices faced by listeners into two categories, in which mental predictions may (P+) or may not (P-) influence referential interpretation in the task setting (Fig. 1C; see also Fig. S4 for trial type definition, simulation, and test statistics) (*22*).

Consistent with previous research (*19, 20*), the RSA model explained the listener data well (Fig. 2A), correctly predicting 68.76 ± 0.99% of listener choices in out-of-sample tests (chance level = 33.3%), and outperformed a variety of alternative models (Fig. S3) (*22*). Using the RSA parameter calibrated on listener data, we derived pragmatic likelihood estimates for each listener on each trial. Important for interpreting neuroimaging data, the average pragmatic likelihood estimates matched the aggregate choice patterns of speakers (Fig. 2B), indicating that the pragmatic likelihood estimates reflect not only what speakers should rationally select to achieve a communicative goal but also what they actually selected in the experiment.

**Fig. 2.**
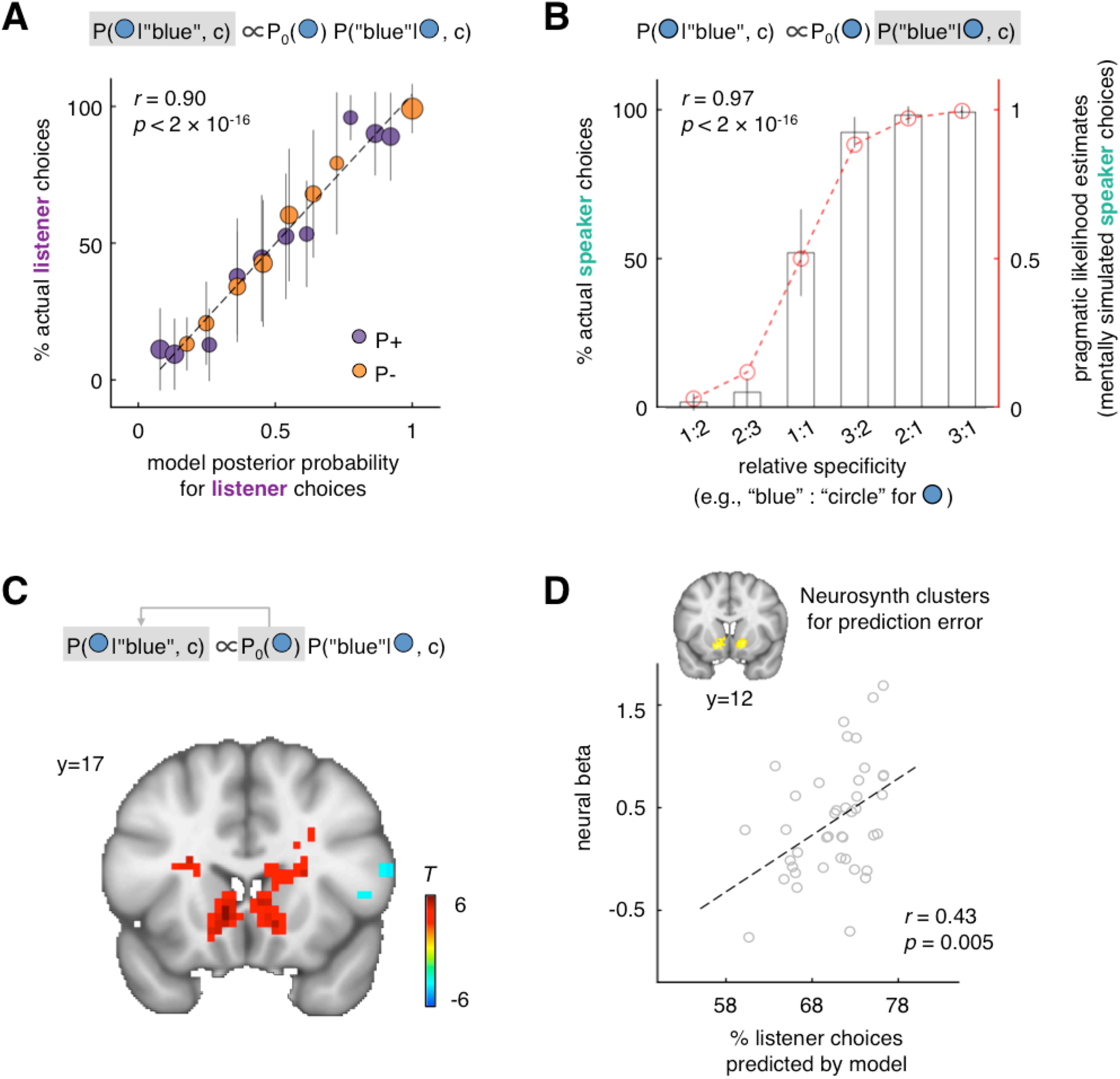
Method validation. **(A)** Actual listener choices conditional on the received referring expression and context as a function of the model-derived posterior probabilities. Data are pooled over all listeners in the symmetric condition and binned by a step size of 0.1 based on model predictions; for single listener results see (*22*). The dashed line represents a perfect model fit. The size of a circle is proportional to the number of observations. **(B)** Actual speaker choices conditional on the target and context (grey bar) matched the pragmatic likelihood estimates derived from the listener data (red dashed line). According to RSA, the pragmatic likelihood estimate for a possible target is a softmax function of the relative specificity between the competing expressions associated with the object (x-axis) (*22*). **(C)** Consistent with previous studies (*23, 24*), the listener bilateral striatum encodes the update signal (posterior – prior probability) for the chosen object at the time of expression onset [*P* < 0.05 cluster-wise family-wise error rate (FWE) corrected, cluster-forming threshold *P* < 0.001; see also Table S1]. **(D)** Across listeners, a superior RSA model fit to the listener data is associated with enhanced neural responses to the update signal in an independent region of interest (ROI) for learning and updating (using the term “prediction error”) defined from Neurosynth (*25*), which is predominately confined to the nucleus accumbens. Each circle represents a listener.

Next, we investigated whether the listener’s brain activity reflected key computational components derived from RSA, including the Bayesian update and pragmatic likelihood signals, at the point when listeners received messages from speakers. A standard general linear model (GLM) analysis revealed that the Bayesian update signal, as assessed by the difference between the prior and model-derived posterior probability that the chosen object was the intended referent, scaled with activity in the listener bilateral striatum on a trial-by-trial basis (Fig. 2C; see also Materials and Methods for how prior probability was assessed). This effect is consistent with previous evidence of reward learning and Bayesian updating in decision neuroscience (*23, 24*). In addition, the striatal activity in response to the Bayesian update signal varied across listeners, such that those whose choices were better characterized by RSA showed a greater update-related signal in a region-of-interest (ROI) independently defined for learning and updating by an automated online meta-analysis (Fig. 2D) (*25*).

To look for listener brain regions where a significant amount of variances could be uniquely explained by pragmatic likelihood estimates while controlling for other potentially related variables, we entered the trial type (P+/P-), posterior probability prediction, and pragmatic likelihood estimate as parametric modulators into a single regression model (*22*). This revealed a significant effect of pragmatic likelihood estimates associated with the chosen object in a cluster in the listener ventromedial prefrontal cortex (vmPFC) (Figs. 3 and S7). Additional analyses suggested that the observed vmPFC activity could not be attributed to correlations with Bayesian computational components other than pragmatic likelihood (e.g., prior, posterior, or update) or cognitive factors that the listener’s brain likely encoded during the inferential process (e.g., message type, context configuration, reaction time, and choices). The reported effect remained robust to the inclusion of all these variables as regressors of no interest into a single regression (Figs. S6 and S8-9), and were predictive of the speakers’ actual choices above and beyond the model-derived pragmatic likelihood estimates (Fig. S10).

**Fig. 3.**
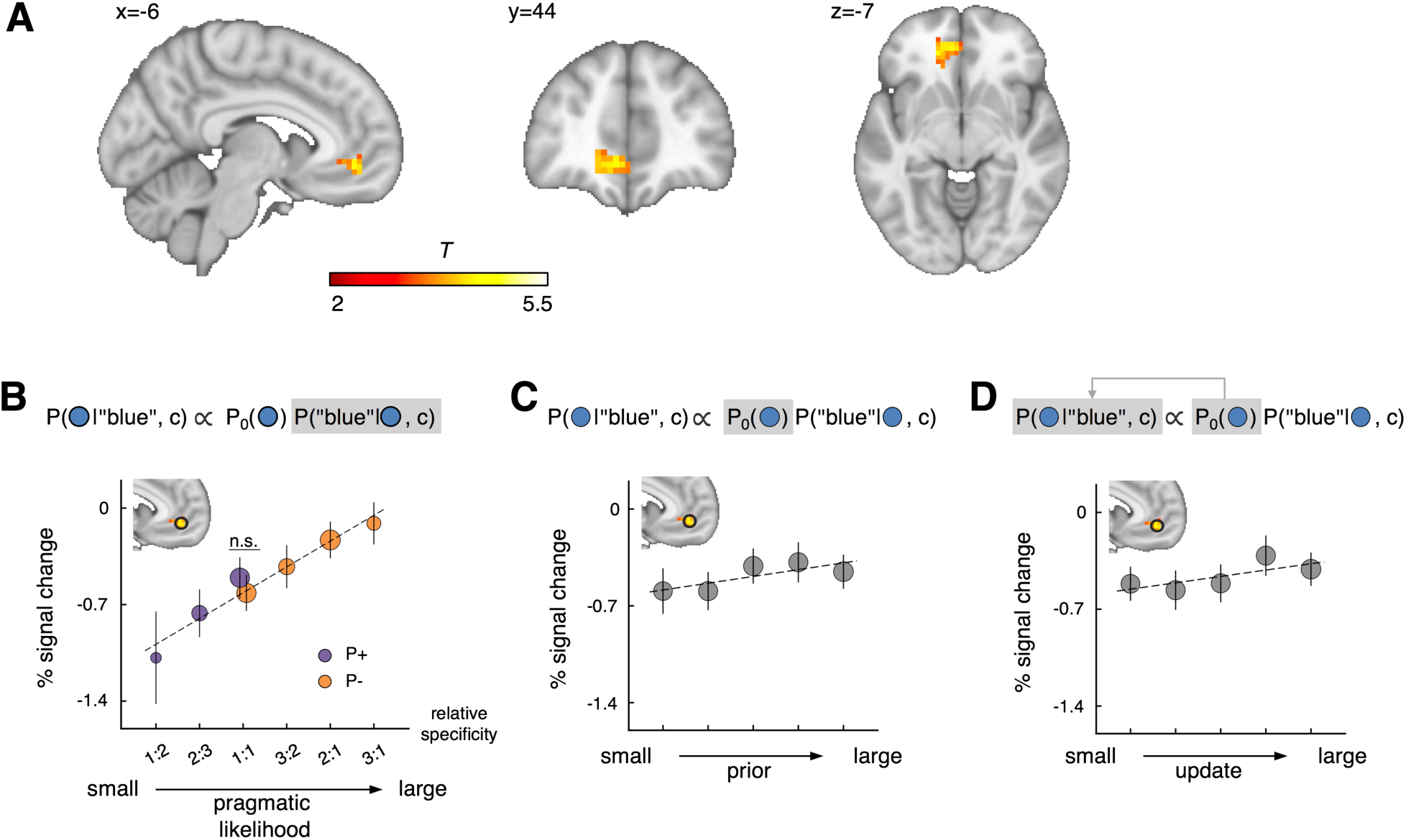
Listener vmPFC encodes pragmatic likelihood estimates. **(A)** Significant activation of the listener vmPFC with respect to pragmatic likelihood estimates for the chosen object upon receiving the speaker’s message in the symmetric condition (*P* < 0.05 cluster-wise FWE corrected, cluster-forming threshold *P* < 0.001). **(B)** BOLD signals extracted from the vmPFC ROI (y-axis), defined as a 6-mm ball around the peak voxel (MNI: -6/44/-7), against the pragmatic likelihood estimates ranked by the relative specificity of the chosen object (x-axis). A mixed-effect linear regression shows the regression coefficient for the vmPFC ROI signal against pragmatic likelihood estimate is 0.67 ± 0.14 (*t*_40._ = 4.68, *P* = 9.8 × 10^−5^; Bonferroni corrected). Note that, by design, P- trials are generally associated with higher pragmatic likelihood estimates relative to the P+ trials, but the listener vmPFC tracks pragmatic likelihood estimates in both P+ and P- trials with no significant difference in the slopes (Δ*β*= 0.01 ± 0.42, *t*_40_ = 0.02, *P* = 0.982), demonstrating similar responses when pragmatic likelihood estimates are equal at the relative specificity 1:1 (P+ = −0.50 ± 0.15 vs. P-= −0.62 ± 0.13, *t*_40_ = 0.94, *P* = 0.354). See also Fig. S6C for the whole-brain analysis using P- trials only. **(C-D)** BOLD signals extracted from the same vmPFC ROI against the prior probability **(C)** and update signal **(D)** of the chosen object (mixed-effect linear regression coefficient, prior = 0.33 ± 0.20, *t*_40._ = 1.63, *P* = 0.333; update = 0.33 ± 0.18, *t*_40._ = 1.77, *P* = 0.252 ; all Bonferroni corrected). For whole-brain analyses of the prior and update signals see Fig. 2C and Fig. S9. Error bars represent inter-subject SEM. Circle sizes represent sample sizes.

Strikingly, the pragmatic likelihood estimate was encoded in the listener vmPFC, and only vmPFC, even when not required for decoding speaker intentions. A separate whole-brain GLM regression showed that within P- trials, activity in an overlapping vmPFC cluster was strongly correlated with pragmatic likelihood estimates of the chosen object, with an effect size similar to that in the P+ trials (Fig. S6C, top). Moreover, signals extracted from the listener vmPFC ROI demonstrated similar level of activation in the P+ and P- trials that were associated with the same pragmatic likelihood estimate values (Fig. 3B, at the relative specificity 1:1). These findings suggest an automatic neural simulation of speaker behavior, even when such simulation is irrelevant for utterance interpretation.

These results thus raise the question of what neural systems inform or facilitate the mental simulation signals observed in the vmPFC. Based on previous studies (*8*), we hypothesized that the pragmatic likelihood computation likely involves the communication between the listener’s vmPFC and mentalization network. In line with this hypothesis, psychophysiological interaction (PPI) analyses showed functional coupling—varying according to the RSA model predictions— between a listener’s vmPFC and brain regions typically implicated in mentalization (*22*), including the dorsomedial prefrontal cortex and temporoparietal junction (*26*) (Fig. S11).

To what extent does the vmPFC activity reflect the mental simulation of speakers in other communicative situations? Results from two additional experimental conditions suggest that, whereas varying communicative stimuli did not affect the vmPFC encoding (*symmetric-garment condition*, Fig. 4, A-C and Fig. S6) (*22*), altering the knowledge shared between communicators significantly perturbed the vmPFC signal (a*symmetric condition*, Fig. 4D-F and Fig. S6). In the latter, we performed the experiment in the same group of listeners, with one important difference: instead of three geometric objects in each trial, speakers were able to see only the target when selecting references. Importantly, listeners underwent the same reasoning task inside the fMRI as in the other conditions but were told that speakers faced only the target during their decisions (Fig. 4D) (*22*). If a listener models the utterance selection process from the speaker’s perspective, then the listener would expect a speaker with a restricted perspective to choose between candidate expressions randomly, regardless of context. This implies that a listener should demonstrate flattened vmPFC activation in the asymmetric condition relative to the symmetric condition. As predicted, we found that listeners were sensitive to the manipulation of common ground, such that now only 66.92 ± 0.15% of targets were correctly identified by listeners, a success rate similar to that of listeners choosing literally in response to random speakers (literal recovery rate = 66.88%; *t*_40_= 0.88, *P* = 0.384) (Fig. S5). Importantly, the listener vmPFC showed blunted responses to the pragmatic likelihood estimates derived from the matching symmetric condition, either at the whole-brain level or within the ROI obtained in the original symmetric condition (Fig. 4, E-F, see also Figs. S12-13 for within- and between-listener comparisons).

**Fig. 4.**
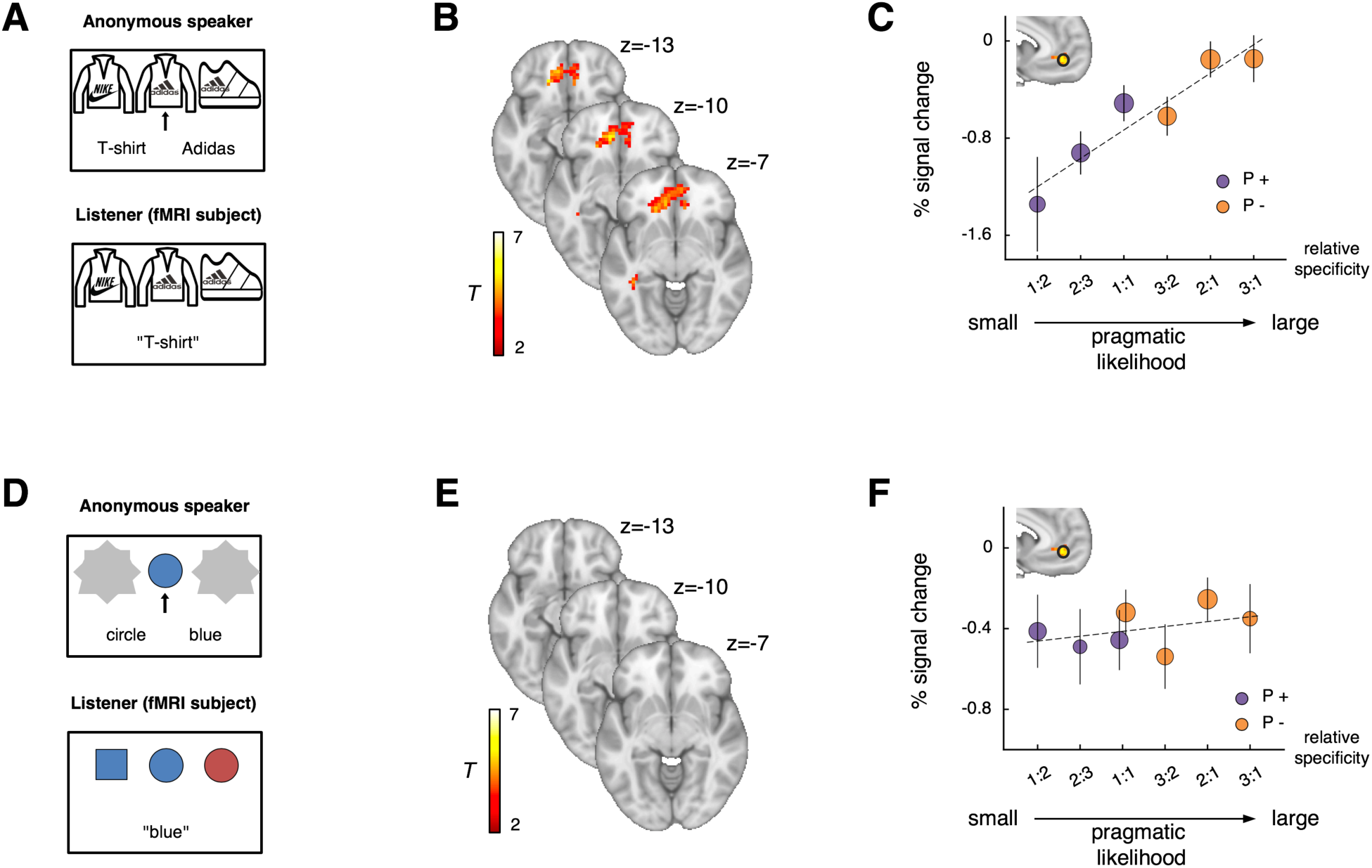
Generality and sensitivity of the pragmatic likelihood representation in the listener vmPFC. **(A)** Schematic of *symmetric-garment condition*, designed for testing whether the vmPFC encoding depends on the specific eliciting stimuli. Different from the other two conditions, the symmetric-garment condition contained less number of trials and only the P+ type at the relative specificity 1:1 (*22*). Both the whole-brain **(B)** and ROI **(C)** analyses revealed a significant correlation between listener vmPFC activity and the pragmatic likelihood estimates of the chosen object (whole brain: *P* < 0.05 cluster-wise FWE corrected, cluster-forming threshold *P* < 0.001, only positive activation was presented for confirmatory purpose, see also Fig. S6 for full activation maps; ROI:*β* = 0.73 ± 0.17, *t*_40_ = 4.44, 2 = 7.0 × 10^−5^). **(D)** Schematic of *asymmetric condition*, designed for testing whether shrinking common ground between communicators dampens the vmPFC responses to pragmatic likelihood estimates. This condition contained the same set of decision contexts for listeners as in the symmetric condition (*22*). Both the whole-brain **(E)** and ROI **(F)** analyses revealed a diminished correlation between vmPFC activity and pragmatic likelihood estimates derived from the matching symmetric condition (whole brain: 2 < 0.05 cluster-wise FWE corrected for positive activation, cluster-forming threshold *P* < 0.001; ROI: *β*= 0.19 ± 0.14, *t*_40._ = 1.38, *P* = 0.177). The same vmPFC ROI was used as in the symmetric condition. Error bars represent inter-subject SEM.

Dating back to Grice’s cooperative principle (*1*), understanding what is meant from what is said in context is thought to involve an inferential process guided by the expectation that the speaker communicates cooperatively. The consistent results across three experimental conditions provide substantial evidence that the listener vmPFC encodes mental simulations of the speaker choice process, complied with rational cooperative principles, inferred from a specific context and common ground information, independent of eliciting stimuli, and irrespective of whether such a simulation is required for utterance interpretation. The rational simulation signal in vmPFC is likely supported by inputs from the mentalization network. Importantly, the finding that the vmPFC signal resembles a Bayesian likelihood function, together with the fact that the listener’s striatal activity correlates with the update from the Bayesian prior to posterior probability, support a mechanism by which the frontal-striatal circuits are engaged in building and then inverting a choice model of the speaker to produce pragmatic interpretations, in a manner mimicking Bayesian inferences.

The result that vmPFC encodes rational simulations of speakers corroborates previous findings that vmPFC is involved in calibrating social actions by processing implied, rather than explicit, social information (*27, 28*). Our data extend these past findings by characterizing the exact role of vmPFC in encoding the speaker’s intention-action contingency, inferred from a specific communicative context, and by specifying the automaticity and generality of the vmPFC representation. This result also supports the view that vmPFC contributes to representing “cognitive maps” that organize task-relevant components for decision-making, especially when behaviors require flexible evaluation of cues, contexts, and actions (*29, 30*), similar to the case of communication.

The finding that altering common ground information modifies the vmPFC signal provides neural evidence for the idea that knowledge and beliefs shared between communicators critically shape pragmatic inferences (*4, 6*). In addition, it raises an intriguing possibility that additional neurocognitive processes are involved in selecting mental models appropriate to the information structure within communicators, perhaps through hierarchical generative mechanisms (*11*-*13*).

The notion of probabilistic generative model has deep roots in artificial intelligence (*14*) and cognitive neuroscience (*11*) such as visual perception (*15*) and motor control (*31*), and has been recently proposed to account for various social inferences (*10, 12, 13, 16, 17*). Our results shed light for the neural instantiation of this generative account in high-level cognitive systems involved in communication and social reasoning. It also helps to explain why people coordinate and cooperate with strangers in the novel, one-shot situations. Past research on cooperation typically focuses on how the brain anticipates partners’ choices by learning from direct experience, such as repeatedly interacting with the same partner within the same decision context (*32*-*34*). In contrast, our results suggest a neural system for simulating another’s behavior based on rational principles that may substitute for learned expectations, consistent with psychological and economic theories regarding the role of strategic mentalization in a range of mutually beneficial behavior (*18, 35*).

More broadly, by highlighting the utility of connecting tools and ideas from decision neuroscience and those of experimental and computational pragmatics, our study raises exciting questions regarding the degree to which neurocognitive substrates of social decision-making are shared by communication and language (*8, 36, 37*), as well as whether behaviors such as detecting sarcasm or interpreting humor can be modeled as strategic, cooperative choices in the brain and brain-inspired artificial intelligence (*38*).

## Acknowledgments

We thank Chunfang Yan for assistance with data collection.

## Author contributions

Q.M. and L.Z. designed the study. Q.M. conducted the experiments. All authors analyzed the data and wrote the manuscript.

## Data and materials availability

Model code, behavioral and imaging data are available at Open Science Framework.

## Materials and Methods

### fMRI participants

A total of 46 healthy, right-handed volunteers [26 females; age = 20.2 ±1.32 years (mean ± SD)] were recruited for the fMRI experiment from the Neuroeconomics Lab subject pool at Peking University, China. All participants reported having normal or corrected-to-normal eye vision, no colorblindness, and no history of neurological or psychiatric illnesses. Five subjects were excluded from data analyses due to excessive motion (N = 4) and a technical problem in stimuli display (N = 1). Informed consent was obtained by the Ethics Committee at Peking University, China.

### Experimental procedure

Subjects participated in a Referential Game adapted from previous studies (*7, 19, 20*). We first conducted a behavioral session in which 60 subjects participated in the Referential Game in the role of *speakers*. Subjects in neuroimaging sessions subsequently played the role of *listeners* with speakers under a random matching protocol. That is, a listener and a speaker were matched pseudo-randomly at the beginning of each round. The listener received a referring expression previously selected by the speaker and needed to recover the intended referent from the received expression. The random matching between speakers and listeners ensured that the probability of repeated interaction was small, thereby preventing communicators from developing hierarchical mental models to collude with their partners.

No feedback was provided to either communicator during the experiment. That is, speakers did not know which items listeners selected in response to the referring expressions, and listeners did not know whether their choices of referents were correct after each decision.

Before the experiment, all subjects (listeners and speakers) were given instructions, completed a quiz, and performed 3 practice trials to ensure comprehension of the game. Subjects were informed that both communicators would be rewarded if a referent was successfully recovered by the listener in a trial. Subjects were paid at the end of the study, based on the total payoff of 100 randomly chosen trials, plus a show-up fee (150 CNY for fMRI listeners and 40 CNY for speakers).

### Experimental conditions

The fMRI experiment included three conditions: symmetric, asymmetric, and symmetric-garment conditions. In the *symmetric condition* (152 trials divided into 2 scanning sessions), a set of three geometric objects were presented to both speakers and listeners in each trial and were known to be presented to both. The speaker was additionally presented with an arrow, randomly distributed among three displayed items, indicating the target object that the speaker needed to refer to and the listener needed to recover based on the received expression. In the *asymmetric condition* (152 trials divided into 2 scanning sessions), we reduced the common knowledge shared between the communicators such that speakers saw only the target object, whereas listeners were informed of all three objects and the fact that speakers were able to see only the target. We also included a *symmetric-garment condition* (72 trials, 1 scanning session) only for listeners as a robustness check for whether the main neuroimaging result depended on specific eliciting stimuli.

During scanning, the symmetric and asymmetric conditions were presented in a block-wise manner with a counterbalanced order [i.e., two successive sessions of the symmetric condition followed by (or following) two sessions of the asymmetric condition]. The symmetric-garment session was always administered at the end. Within each scanning session, trial order was randomly shuffled with a unique order per listener.

### Experimental stimuli

A schematic representation of the Referential Game and the timeline of the experiment is shown in Fig. S1.

**Fig. S1.**
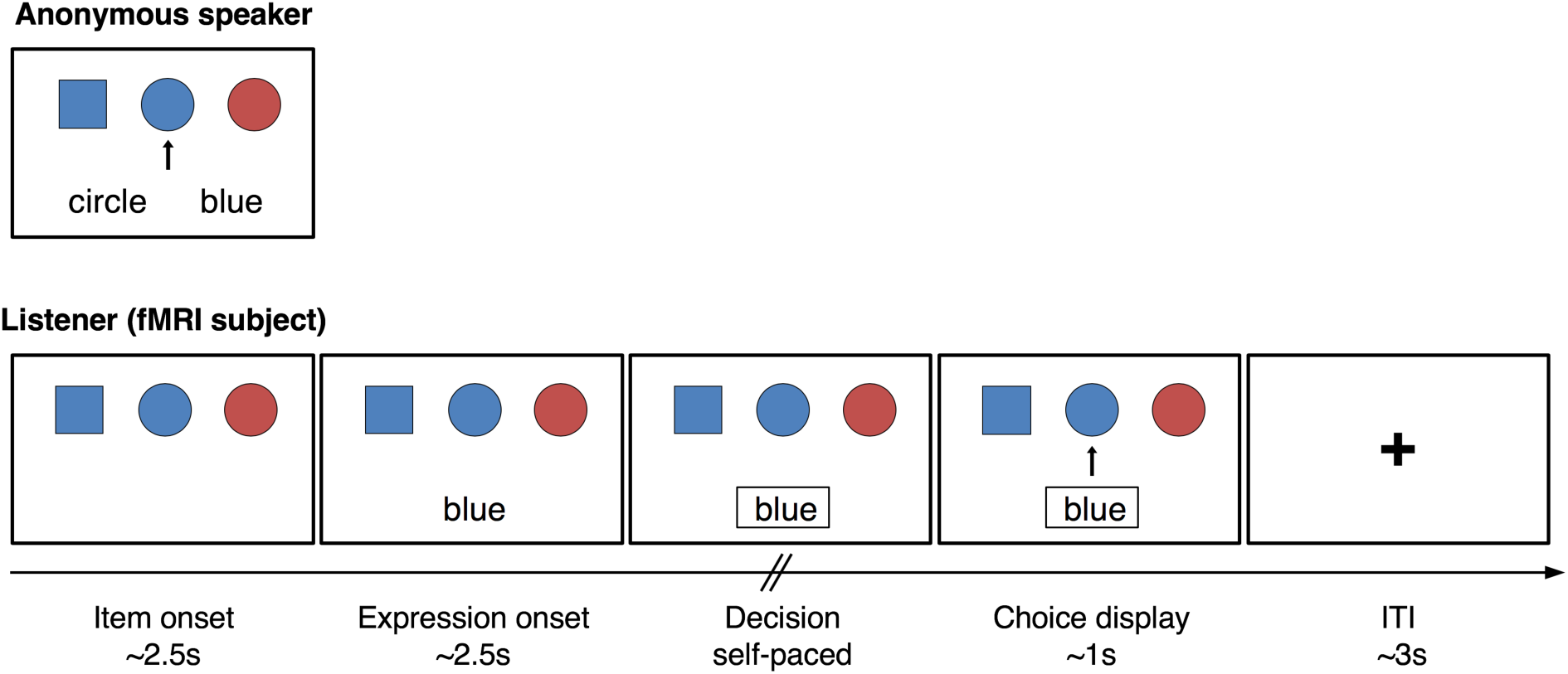
Schematic representation of the symmetric condition. A listener and a speaker are presented with the same context consisting of 3 items in the symmetric condition. The speaker needs to refer to the target item (indicated by an arrow) using either the color or shape of the target, whereas the listener needs to recover the target referent based on the expression sent by the speaker. More specifically, on each trial, a listener is presented with a set of three items for an average of 2.5 s. The referring expression chosen by the speaker is then shown on the screen for an average of 2.5 s. The listener is able to make a self-paced decision once the received expression is framed by pressing buttons mapped to item locations on a response pad. Once the listener makes the choice, an arrow is shown underneath the chosen item for an average of 1 s, followed by a fixation screen for an average of 3 s.

On each scanning trial in the symmetric and asymmetric conditions, a listener is presented with a new *context* consisting of 3 geometric items displayed horizontally, with item locations fixed for the listener and the speaker within each context. All listeners faced the same 304 contexts that contained a total of 16 different geometric objects, generated from 4 colors (red, green, blue, and yellow) and 4 shapes (diamond, square, circle, and trapezoid). All color/shape features can be denoted by a two-character noun in Chinese. We constructed the 304 contexts pseudo-randomly by drawing 3 items out of 16, with replacement, without distinguishing between drawing orders or item locations (Fig. S2). A full list of experimental stimuli is included in the supplemental data (Data S1). The target location was randomly distributed. In the asymmetric condition, only the target was revealed to the speaker, whereas the two distractors were covered by grey masks.

**Fig. S2.**
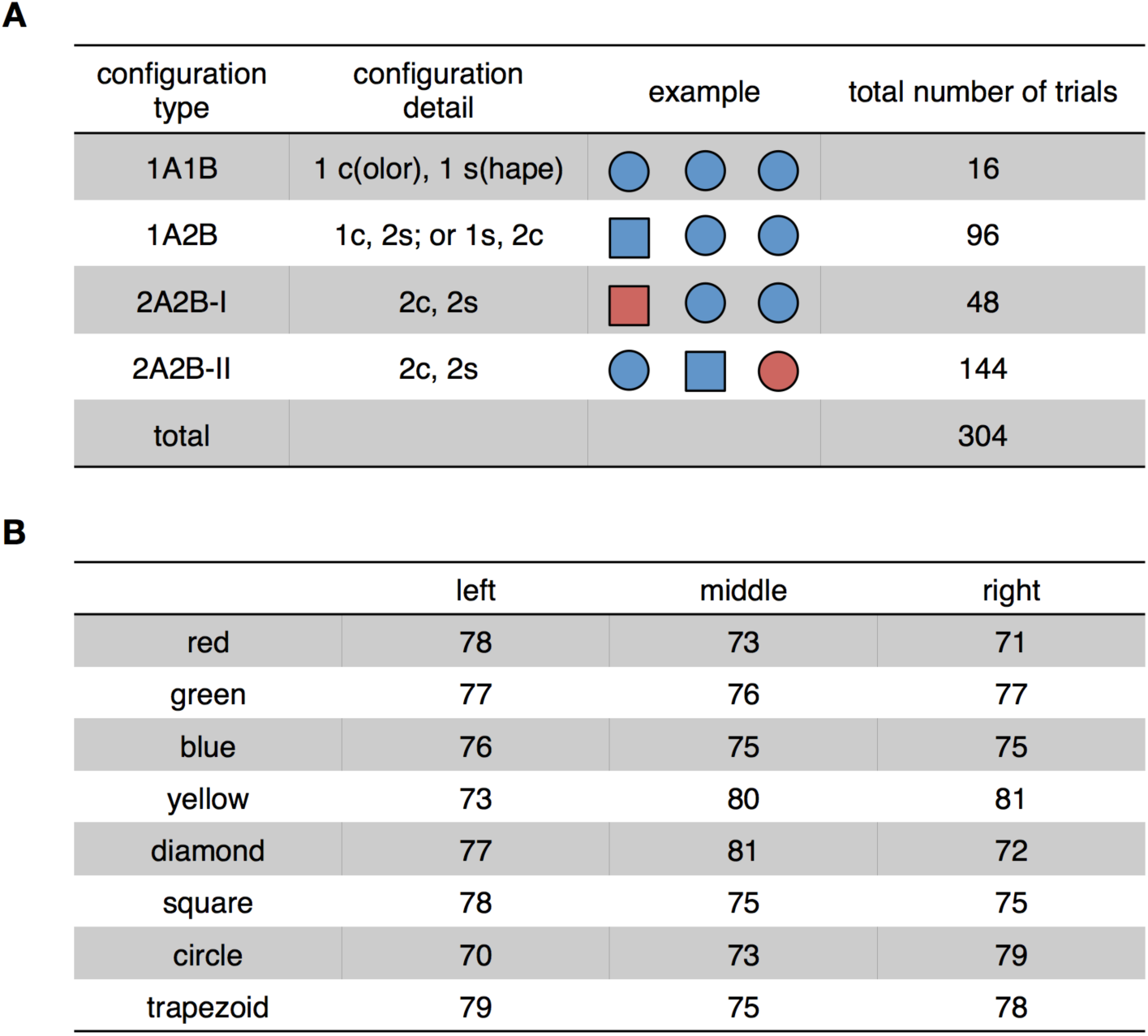
**(A)** Context configuration. Contexts were constructed by generating all possible combinations of 3 objects, randomly drawn from 16 (i.e., 4 colors × 4 shapes), with replacement, that did not distinguish between drawing orders or item locations. We included all these combinations in the experiment with two exceptions, due to the fMRI experiment’s time constraint (1.5-hour scanning time under the current design). We excluded contexts that contained 3 different colors or shapes and manually selected a subset of contexts for type 2A2B-I (48 out of 144) that displayed no obvious patterns in the distribution of item features. This resulted in 304 contexts of 4 configuration types in the symmetric and asymmetric conditions combined. **(B)** Item features are evenly distributed across item locations in 304 contexts [*χ*^2^ (4) = 2.15, 2 = 0.999].

Stimuli in the symmetric-garment condition were created in a similar fashion and included 9 items, generated by 3 garment types (top, pants, and sneakers) and 3 brand names (Adidas, Nike, and Li Ning). Each of these features was associated with a two-character Chinese noun (Data S1).

### Prior probability evaluation

Following previous studies (*19, 20*), we empirically measured the prior probability distribution of target items in 304 contexts using online surveys (https://www.wjx.cn) in a separate sample of Chinese participants (N=900). In each trial, survey participants were presented with 3 geometric items and asked to infer the referent based on an unknown expression in a foreign language. We instructed subjects to follow their intuition and make a guess if they did not know the meaning of the expression. Answers from these participants therefore likely reflect the relative visual saliency among geometric items presented in the same context. To monitor the performances of online participants, 32 sanity check questions were included and evenly distributed throughout the survey, where subjects needed to identify the referent based on a Chinese referring expression that uniquely denoted an item in the context. Ninety-eight survey participants who answered incorrectly on more than 30% of sanity check questions were excluded from the data analysis, whereas the rest of the participants answered sanity check questions with an accuracy rate of 91.63 ± 0.32%.

We calculated the trial-wise prior probability distribution of targets by averaging the choices of each item within each context across survey participants. The empirically measured prior probabilities were subsequently used for fitting listener choices in the symmetric condition and imaging data analyses. No prior probability data were collected for the asymmetric or symmetric-garment conditions.

### Computational modeling

We applied the Rational Speech Act (RSA) model (*10, 19*) to characterize the listener behavior observed in the symmetric condition. Listeners make their decisions based on a Bayesian inferential process that can be formalized as

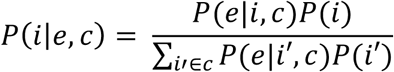

where *P*(*i*|*e,c*) is the posterior probability of a listener choosing a particular item *i* upon receiving an expression *e* in a context *c*; *P*(*e*|*i,c*) is the likelihood that the speaker selects the expression *e* in order to refer to an item *i* in a context *c*; and *P*(*i*) is the prior probability that an item *i* is the target referent. According to the Bayesian setup, listeners need to predict how speakers generate their choices for each possible target in a context in the form of conditional probability distributions *P*(*e*|*i,c*), which we refer to as *pragmatic likelihood*.

RSA assumes that pragmatic likelihood is computed by simulating speaker choices through a rational, goal-directed decision-making model. Specifically, listeners expect that when selecting referring expressions, speakers choose an expression to help the recipient recover the target. In the symmetric condition, this corresponds to choosing the maximally specific (informative) reference within a given context, which can be quantified using an information-theoretic measure, self-information, 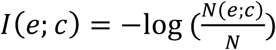, where *N* denotes the total number of items contained in a context *c* (thus, *N* = 3 in our experimental setting), and *N*(*e;c*) denotes the number of objects that an expression *e* can denote in context *c*. For example, if an expression *e* can describe all three items in a context [i.e., *N*(*e;c*) = 3], *e* is not informative at all [i.e., *I*(*e;c*) = 0] and will not help the listener narrow down possible referents.

To convert self-information of candidate expressions into choice probabilities, the model assumes that speaker choices follow a logit or softmax formula widely used in decision-making research:

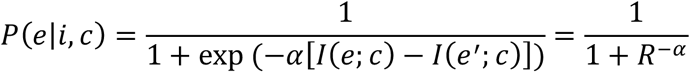

where 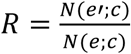 reflects the relative specificity between the expression *e* and its alternative *e*′, and *α* reflects how sensitive speaker choice probability is to the relative specificity between competing expressions, or the “inverse temperature” of the softmax function (e.g., *α* = 0 means listeners expect speakers to select randomly between *e* and *e*′). For example, if an expression *e* is more specific than its alternative *e*′ in referring to a target *i* (i.e., expression *e* can denote fewer items than *e*′), a rational cooperative speaker should be more likely to select *e* over *e*′ [i.e., *P*(*e*|*i,c*) ≥ 0.5]. That is, pragmatic likelihood *P*(*e*|*i,c*) is a non-decreasing function with respect to the relative specificity *R*, as demonstrated in Fig. 1D.

### Model estimation

To calibrate the RSA parameter *α* with listener behavior observed in the symmetric condition, we estimated the behavioral model using both pooled estimation and hierarchical Bayesian analysis. For pooled estimation, we assumed that the choices of all listeners were generated by a single, shared *α*, and we applied the maximum likelihood estimation with grid search over a large non-negative domain for *α*, since the likelihood function may be not globally concave. Specifically, we fit listener choice data by maximizing the log of posterior probability of observed listener choices, ∑_*k*_∑_*t*_log *P*(*i*_*k,t*_| *e*_*k,t*_, *c*_*k,t*_), pooled over listeners *k* and trials *t*.

Second, to account for individual differences in referential interpretation, we also calibrated individual listener parameters using the well-established hierarchical Bayesian model estimation method. We assumed that the parameter *α*for each listener was randomly drawn from a normal distribution governed by group-level mean and variance [i.e., *α*_*k*_ ∼ *N*(*µ,σ*)], whereas the group-level parameters were independently sampled from a uniform prior distribution taking values from 0 to infinity. We computed the posterior likelihood of observing listener choices with the Markov chain Monte Carlo (MCMC) method implemented in RStan (*39*). Two MCMC chains were simulated with 2500 iterations after 2500 burns-in, resulting in 2500 posterior samples for each parameter in each chain. All parameters were checked for convergence both visually (from the trace plot) and through the Gelman-Rubin test (all 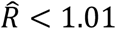).

In the model estimation and subsequent behavioral and neuroimaging analyses, the Bayesian prior probability distribution was empirically measured using an independent online sample. Additionally, we excluded trials in which listeners made literal mistakes for data analyses (e.g., choosing a red circle upon receiving an expression “blue”) to avoid zero probability for the chosen item according to model prediction. This resulted in the removal of 0.50 ± 0.12%, 0.33 ± 0.07%, and 0.58 ± 0.17% trials in the symmetric, asymmetric and symmetric-garment conditions, respectively. According to the pooled estimation result, the best-fitting *α* is 4.97, and the log likelihood of listener choices observed in the symmetric condition is -2351.63. For hierarchical Bayesian analysis, the individual parameter *α*_*k*_∼*N*(5.93, 2.17), and the deviance information criterion (DIC) is 107.36 ± 2.93.

### Model comparison

To further verify the plausibility of RSA and test for alternative decision strategies that may be employed by listeners, we compared RSA with the following models representing competing hypotheses regarding how listeners recognize speaker intentions.

#### Literal listener model

This model assumes that listeners interpret received expressions literally, randomly choosing among items that the received reference can denote within the context. This model contains no free parameter and serves as a baseline for model comparison.

#### Flat prior model

This model assumes that a flat prior probability distribution is used for the Bayesian inferential process within RSA, serving to test the assumption that empirically measured prior probabilities, a measure of relative saliency between items within a context, contribute to referential interpretation. This model also contains a single parameter *α* as in the original RSA.

#### Sophisticated listener model

This model assumes that speakers think one step further than maximizing reference specificity as proposed by the RSA, taking into account the possible choices by listeners in response to a specificity-maximizing speaker. On the other hand, listeners decode speaker intention based on this sophisticated mental model of speaker behavior. In particular, following the well-established cognitive hierarchy approach (*40*), we assume that sophisticated listeners derive the most probable referents through a Bayesian inferential process that can be characterized as

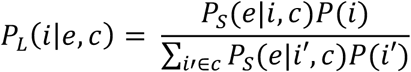

where *P*_*L*_(*i*|*e,c*) is the posterior probability of sophisticated listeners choosing a particular item *i* given the expression *e* and context *c*, and *P*_*S*_(*e*|*i,c*) is the probability of a speaker selecting an expression *e* to refer to *i* in a context *c*. Importantly, and different from the assumption in RSA, here the speaker is assumed to cooperate with an RSA listener according to the following softmax decision rule:

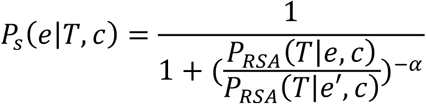

where *P*_*RSA*_(*T*|*e,c*) is the probability of an RSA listener recovering a target *T* from an expression *e* in context *c*, and the so-called RSA listener is a listener who uses Bayesian inferences to derive the target based on the expectation that speakers seek to maximize the specificity of the chosen reference. The sophisticated listener model also contains a single free parameter, *α*, reflecting the choice sensitivity of speakers to the difference between alternative referring expressions.

**Fig. S3.**
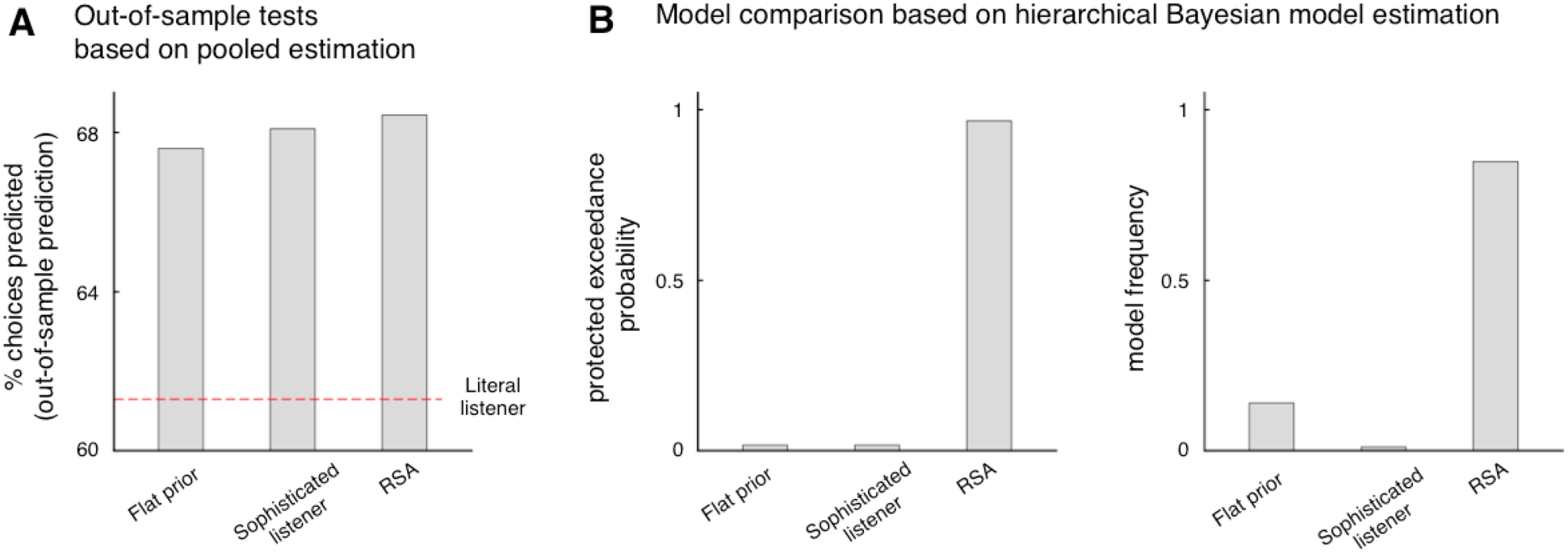
**(A)** Out-of-sample predictive power based on pooled estimation is superior for RSA compared with the alternatives (RSA vs. flat prior: *t*_*40*_ = 4.09, *P* = 0.0004; RSA vs. sophisticated listener: *t*_*40*_ = 3.63, *P* = 0.0016; all Bonferroni corrected). Models are fitted using the pooled estimation method with either the odd-numbered or even-numbered trials in the symmetric condition compounded over listeners. The predicative power is computed by averaging the hold-out prediction accuracy for each model. The dashed line represents the explanatory power of the literal listener model, which contains no free parameter and therefore does not need model fitting. **(B)** We also implement the Bayesian model selection method for models with free parameter, assuming that listeners follow different decision models with a fixed but unknown population distribution of the parameter. Both the protected exceedance probability (left) and model frequency (right) calculated by Bayesian model selection based on DIC (*41*) show superior RSA fit relative to models with alternative hypotheses regarding either the prior probability (flat prior model) or pragmatic likelihood (sophisticated listener model).

We compared the fits of listener choices among competing models and found the highest predictive power by RSA using either pooled estimation or Bayesian model selection (*42*), according to either in-sample or out-of-sample measurements of goodness of fit (Fig. S3).

### Prediction+ (P+) and prediction-(P-) trials

We classified choices faced by listeners in the symmetric condition into two categories, depending on whether predicting speaker choices is critical for referential interpretation. In particular,

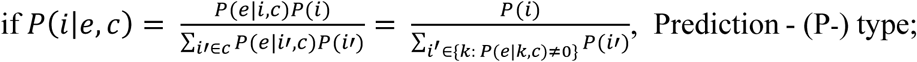

otherwise, Prediction+ (P+) type.

Put in other words, the posterior probability distribution that an object would be referred to depends only on the prior probability but not the pragmatic likelihood in the P- trials, suggesting that listeners can arrive at the same referential interpretations in these situations without simulating the speaker’s choice behavior. More precisely, under the current experimental setup, the P-type consists of trials where a received reference denotes either a single item or a number of identical items (i.e., same color and shape) in the context.

**Fig. S4.**
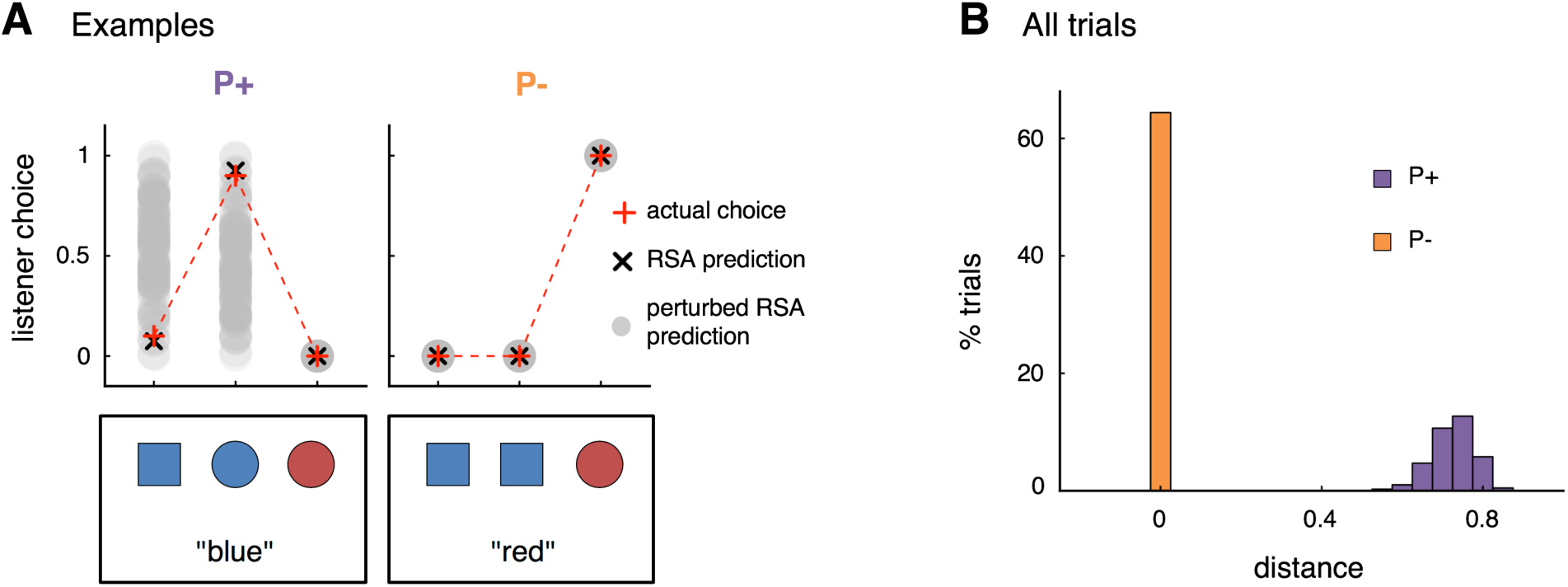
**(A)** P+ and P-examples show that perturbing pragmatic likelihood influences RSA model predictions in the P+ trials but not P- trials. Red crosses represent the actual choice frequencies of listeners in the corresponding decision in the symmetric condition. Black crosses represent the posterior probability distribution of listener choices derived from the best-fitting, group-level RSA estimation. Grey dots are the posterior probabilities simulated from RSA based on 100 randomly perturbed pragmatic likelihood values, in which randomly generated probability distributions are assigned as the pragmatic likelihood over items that can be literally described by the received references. The distributions of grey dots are similar for the blue square and the blue circle in the left example, because both items can be described as “blue” and are both assigned with perturbed pragmatic likelihood randomly. **(B)** Classification outcomes across trials in the symmetric condition. Histograms depict the average Euclidean distance between the posterior probability predictions generated by the best-fitting RSA and 100 randomly perturbed RSA.

To verify and visualize classification outcomes, we generated RSA model predictions using randomly perturbed pragmatic likelihood values and compared these predictions with the actual listener choices in the symmetric condition (Fig. S4). In P+ trials, pragmatic likelihood critically shapes the utterance interpretation such that perturbing pragmatic likelihood gives rise to differential interpretations of the same message. In P- trials, however, listeners always arrive at the same interpretation even when pragmatic likelihood is randomly perturbed, leaving it difficult to determine whether a listener actually generates rational predictions regarding the speaker as proposed by RSA. Neuroimaging data, on the other hand, offer an opportunity for testing the extent to which the mental simulation of speakers is related to pragmatic reasoning in a manner implied by computational models incorporating rational, cooperative assumptions.

### fMRI data acquisition and preprocessing

We collected the fMRI images for each listener using a 3T Siemens Prisma scanner and a 32-channel head coil at the Center for MRI Research at Peking University. Images were acquired using echo-planar T2* images with BOLD (blood-oxygenation-level-dependent) contrast and angled 30° relative to the AC-PC line to minimize susceptibility artifacts in the orbitofrontal area. The scanning parameters are as follows: repetition time (TR) = 2000 ms, echo time (TE) = 30 ms, flip angle = 90°, field of view (FoV) = 192 × 192 mm, slice thickness = 4 mm, slice gap = 0.4 mm, voxel size = 3 × 3 × 4 mm3,32 slices. A high-resolution T1-weighted structural image was acquired using a magnetization-prepared rapid gradient echo sequence (MPRAGE) with the following parameters: TR = 2530 ms, TE = 2.98 ms, flip angle = 7°, FoV = 224 × 256 mm, slice thickness = 1 mm, slice gap = 0.5 mm, voxel size = 0.5 × 0.5 × 1 mm3, 192 slices.

Imaging preprocessing and analyses were performed in SPM12 (https://www.fil.ion.ucl.ac.uk/spm/software/spm12/) with MATLAB R2016b. For each fMRI session, the raw images were first slice-timing corrected and then aligned to the first volume to correct participants’ head motion. After that, the images were spatially normalized into the Montreal Neurological Institute (MNI) template with a final image resolution of 3 × 3 × 3 mm3 and smoothed using a 6-mm FWHM Gaussian kernel. All images were temporally filtered using a high-pass filter with a width of 128 s.

### fMRI data analysis

We implemented a generalized linear model (GLM) for model-based fMRI analysis widely used in the field of decision neuroscience. The best-fitting RSA model parameter from pooled estimation was used to calculate the trial-wise pragmatic likelihood, posterior probability, and posterior–prior update (with prior probability obtained in a separate online sample) for each listener. These values were then used as parametric modulators in the model-based fMRI analysis for listener brain activity observed in the symmetric condition. To examine the robustness of neural encoding of pragmatic likelihood, we also included a single scanning session of the symmetric-garment condition, where we computed the corresponding pragmatic likelihood values for each trial and each listener, assuming listeners in the symmetric-garment condition shared the same *α* estimate as in the symmetric condition. Finally, to test whether altering common ground between communicators modifies the neural encoding of pragmatic likelihood, we included the asymmetric condition, where we entered the same pragmatic likelihood value from the matching symmetric trial as the parametric modulator for fMRI analysis.

In GLMs, each trial was modeled as 4 discrete events—item onset, expression onset, choice submission, and fixation onset—all as stick functions (i.e., duration = 0). Regressors were convolved with the canonical hemodynamic response function and entered into a regression analysis against each listener’s BOLD responses. We were specifically interested in listeners’ brain activity when they received the referring expression from the speaker; thus, variables of interest were entered into GLMs as parametric modulators associated with expression onset.

In particular, the first GLM served to establish the validity of the RSA model at the neural level; thus, it included the posterior–prior update signal associated with the chosen object as the parametric modulator for trials in the symmetric condition (Fig. 2C). In the second GLM, we sought to establish the neural signature of pragmatic likelihood by including the following parametric modulators: trial type (i.e., P+/P-), model-derived posterior probability for the chosen object, and the pragmatic likelihood estimate for the chosen object, with the default automatic orthogonalization switched off (Figs. 3A and S7). To examine the robustness of pragmatic likelihood encoding and control for the influences of potential confounding factors, the third GLM included the following 9 parametric regressors: trial type (P+/P-), posterior probability of the chosen item, message type (color/shape), context configurations (1A1B/1A2B/2A2B), reaction time, choice (left/mid/right), choice uncertainty (entropy of the posterior probability), outcome uncertainty (distance between posterior probability of the chosen object and 0.5), and pragmatic likelihood for the chosen item (Fig. S6B). The fourth GLM served to explore the extent of neural activation in response to pragmatic likelihood and thus included only a single parametric modulator of the pragmatic likelihood estimates for chosen objects with no control variables (Fig. S6A). The 6 vectors of head motion parameters derived from pre-processing were also included as nuisance regressors in all analyses. Regression betas from each listener were averaged across sessions within each condition and then taken into random-effects group-level analyses.

All whole-brain analyses were thresholded and displayed at the family-wise error rate (FWE)-corrected *P* value (*P*_FWE_) of 0.05 at the cluster level, with a cluster-forming threshold of *P*_unc_. < 0.001, as reported by SPM. In addition, similar whole-brain results were obtained with a non-parametric thresholding approach applied to the second-level analyses using default settings in SnPM13 (*43*) (i.e., 5,000 permutations, cluster-forming threshold of 0.001, *P*_FWE_ < 0.05).

### Functional connectivity analyses

To test whether listener vmPFC differentially connected with areas within the well-established theory-of-mind (ToM) network according to behavioral model predictions, we analyzed functional connectivity between listener vmPFC and ROIs that were a priori selected using Neurosynth (http://www.neurosynth.org) for the term “theory of mind”, followed by an exploratory whole-brain analysis to identify areas other than the a priori defined ToM ROIs that show similar effects. The vmPFC cluster identified in Fig. 3A was used as the seed region for PPI analyses. Four ToM ROIs were defined by 6-mm spheres around peaks of the map automatically generated by Neurosynth for “theory of mind” [dmPFC: (4, 58, 24), LTPJ: (-54, -54, 22), RTPJ: (58, -54, 20), and Precuneus: (-2, -56, 40)]. We performed PPI analyses in SPM12 with the following regressors for the event of expression onset: (i) the average BOLD time series extracted from the vmPFC cluster, (ii) the dummy variable indicating whether a listener choice follows the RSA model recommendation (i.e., the choice is assigned with the highest posterior probability by the best-fitting model), and (iii) the interaction term between the average vmPFC time course and the dummy variable.

