## Supplemental Figures and table for "Listener’s vmPFC simulates speaker choices when reading between the lines"

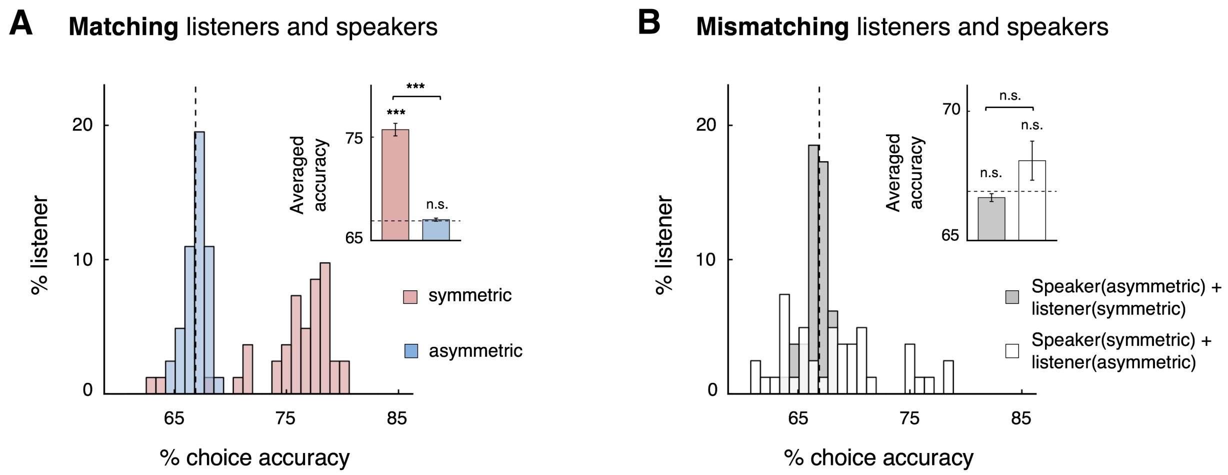


**Fig. S5. (A)** Distributions of recovery rates in referential interpretation in the symmetric (pink) and asymmetric (blue) conditions. Histograms illustrate the percentage of listeners for each bin of recovery accuracy under different conditions. **(B)** Mismatching listeners and speakers across different conditions (e.g., speaker in the asymmetric condition paired with listeners in the symmetric condition) decreases the recovery rates, suggesting that the choices of both listeners and speakers, rather than those of only the speakers or listeners, contribute to the recovery accuracy observed in the symmetric condition. The dashed line represents the choice accuracy associated with literal listeners who always select randomly among all items that can be literally denoted by the received expression. Error bars indicate inter-subject SEM. *$P < 0.05$, **$P < 0.01$, ***$P < 0.001$; n.s. not significant; all Bonferroni corrected.

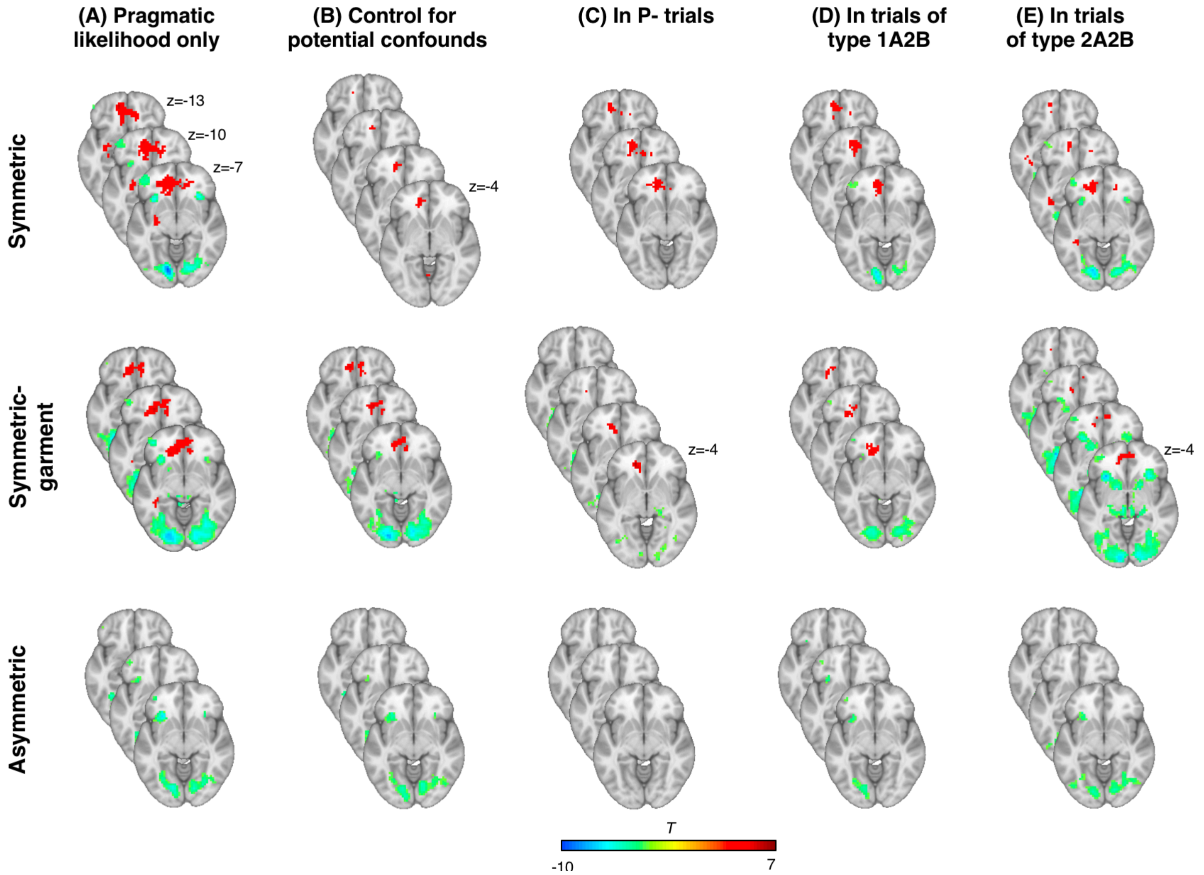


**Fig. S6.** Robustness checks for the vmPFC encoding of pragmatic likelihood estimates. **(A-B)** To demonstrate the extent of vmPFC activation in response to pragmatic likelihood estimates, we performed whole-brain searches for pragmatic likelihood estimates associated with the chosen item, for each condition, with or without controlling for decision variables that the listener's brain likely encodes at the expression onset. The variables of no interest for the regression of the symmetric condition include trial type (P+/P-), posterior probability of the chosen item, message type (color/shape), context configuration (1A1B/1A2B/2A2B), reaction time, choice type (left/middle/right), choice uncertainty (entropy of the posterior probability), and outcome uncertainty (distance between posterior probability of the chosen object and 0.5). For the symmetric-garment and asymmetric conditions, we included all variables above except those related to the model-derived posterior probabilities, because no prior probability data were collected and thus no model estimation was performed for these conditions (see also materials and methods for prior probability data collection). **(C)** To demonstrate that the listener vmPFC tracks pragmatic likelihood estimates even when mental simulation is irrelevant for referential interpretation, we performed a whole-brain search for pragmatic likelihood estimates in the P- trials only. **(D-E)** To demonstrate that listener vmPFC encoding is robust to context configurations, we performed whole-brain search for pragmatic likelihood estimates, in trials of type 1A2B or 2A2B, separately (see also Fig. S2 for trial configurations). All results are thresholded and displayed at cluster-level $P_{\mathrm{FWE}} < 0.05$, with a cluster-forming threshold $P_{\mathrm{unc.}} < 0.001$, except (B), where results are presented at $P_{\mathrm{unc.}} < 0.001$, $k > 20$.

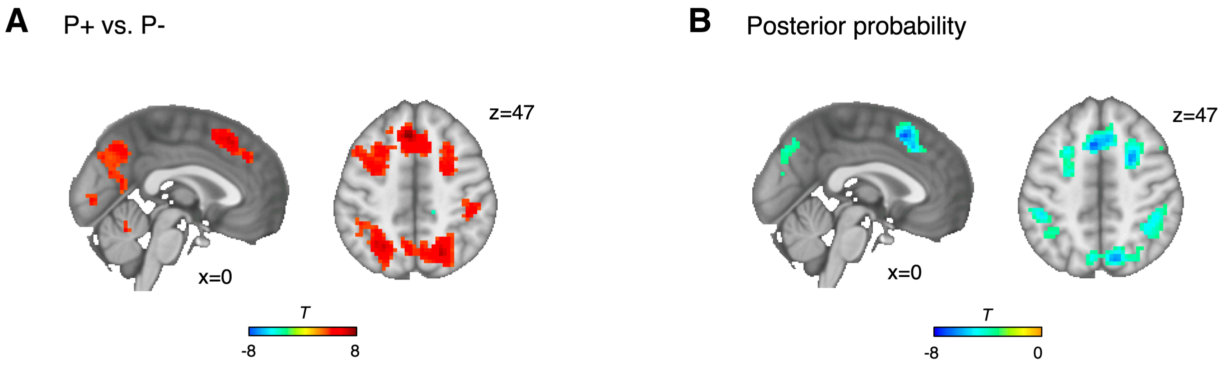


**Fig. S7.** Related to Fig. 3A, brain regions where activity uniquely responds to **(A)** the trial type (P+/P-) or **(B)** model-derived posterior probability estimate of the chosen object, when the trial type, posterior probability, and pragmatic likelihood estimate for the chosen object were entered as parametric modulators into a single GLM with the automatic orthogonalization turned off (see Materials and Methods). All results are thresholded and displayed at cluster-level $P_{\mathrm{FWE}} < 0.05$, with a cluster-forming threshold of $P_{\mathrm{unc.}} < 0.001$.

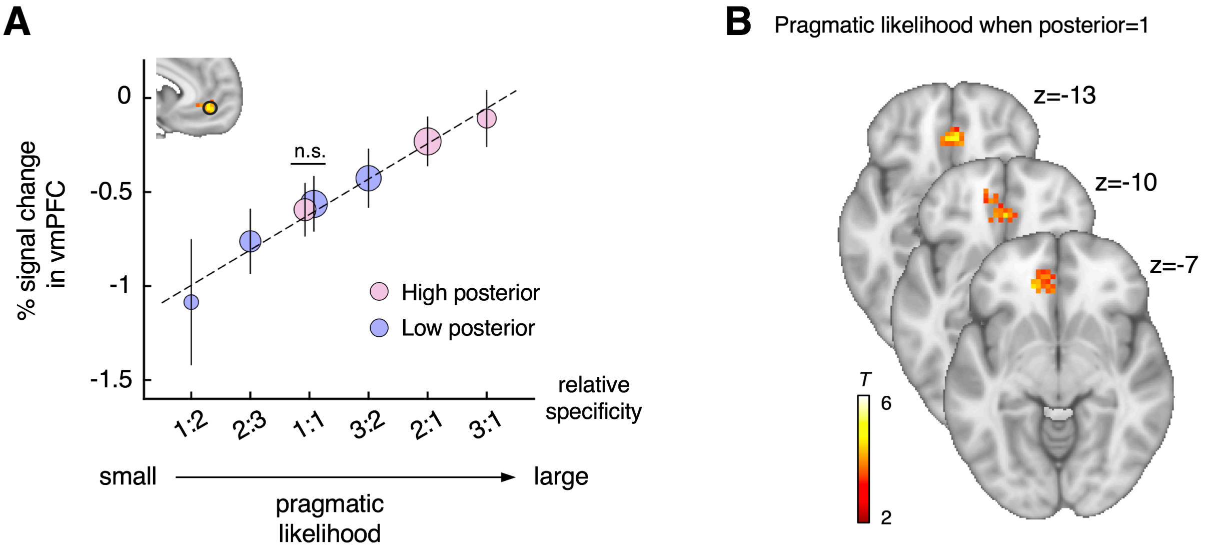


**Fig. S8.** The vmPFC encoding of pragmatic likelihood estimates cannot be attributed to the responses to the posterior probability estimates. **(A)** vmPFC signals against pragmatic likelihood, conditional on model-derived posterior probability for the chosen object. The percent change in the hemodynamic signal was averaged across all voxels identified in Fig. 3A at the onset of referring expression. The means ± SEM of the resulting signal are plotted for each specificity ratio and colored for high (pink) or low (purple) posterior probability estimates of the chosen object based on median splits within each fMRI session of each listener. Error bars indicate inter-subject SEM. **(B)** Whole-brain search results for pragmatic likelihood estimates of the chosen object at expression onset in trials when posterior probability estimates of the chosen object are equal to 1 (thresholded and displayed at cluster-level $P_{\mathrm{FWE}} < 0.05$, with a cluster-forming threshold of $P_{\mathrm{unc.}} < 0.001$). That is, even in trials where the posterior probability values are constant, listener vmPFC activation is still significantly associated with the pragmatic likelihood estimates in the symmetric condition.

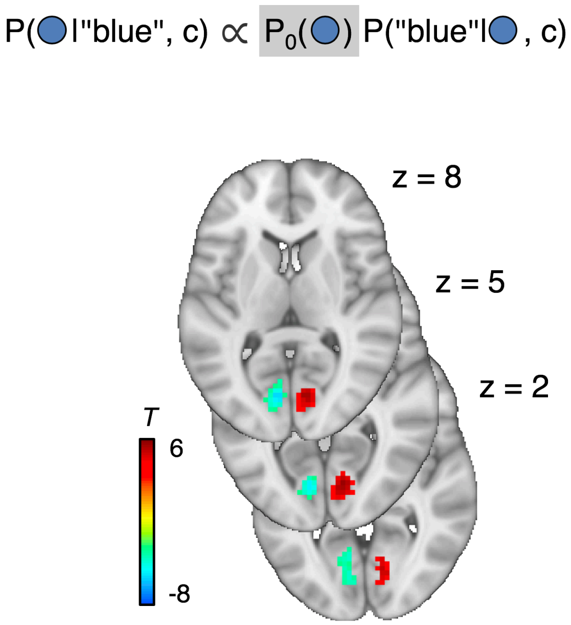


**Fig. S9.** Neural encoding of the prior probability distribution that an object would be referred to at the time of expression onset in the symmetric condition. Consistent with previous evidence (*44*) and the hypothesis that the prior probability distribution reflects the relative visual saliency among objects presented in the same context, activation in the bilateral occipital cortex reflects the location of the object with the highest prior probability in context (left = -1, middle = 0, and right = 1). The prior probability was empirically measured by an online survey in a separate sample (see materials and methods), following previous studies (*19*, *20*). The results are thresholded and displayed at cluster-level $P_{\mathrm{FWE}} < 0.05$, with a cluster-forming threshold of $P_{\mathrm{unc.}} < 0.001$.

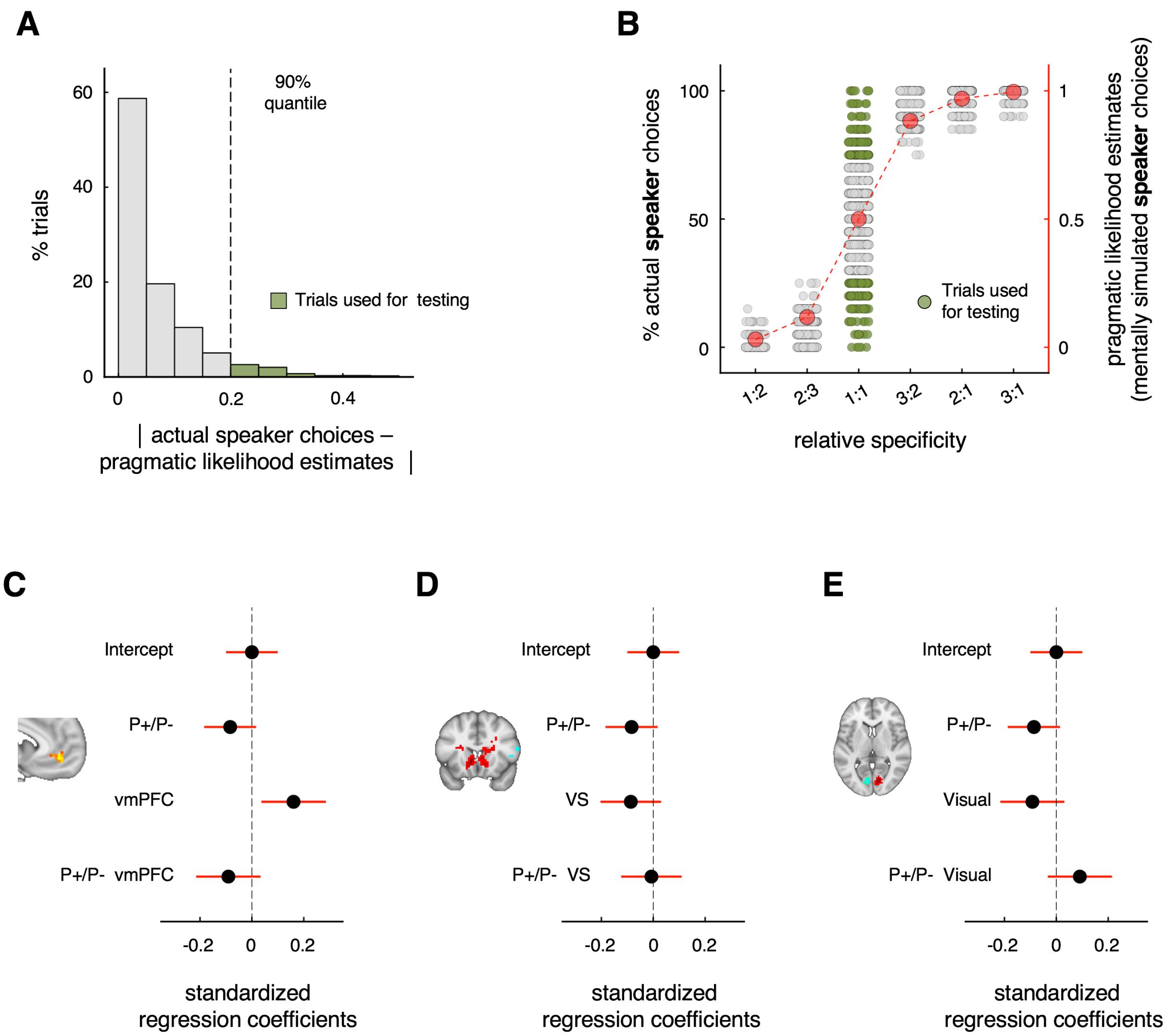


**Fig. S10.** BOLD signals extracted from the listener vmPFC ROI predict the speaker's actual choice patterns, even in trials in which the pragmatic likelihood estimates fail to do so. If the listener vmPFC activity reflects the mental simulations of speakers, and if the speaker choices contain any behavioral nuances that the pragmatic likelihood estimates fail to capture but listeners correctly anticipate during communication, then the listener vmPFC activation should predict the actual choice pattern of speakers above and beyond the pragmatic likelihood estimates derived from the best-fitting RSA model. We tested this hypothesis by examining the degree to which vmPFC activation outperforms the model-derived pragmatic likelihood estimates in predicting the likelihood that speakers chose an expression to refer to a given target in the symmetric condition. To avoid ceiling effects, we sorted trials from all listeners according to how well the pragmatic likelihood estimates could explain speaker behavior and focused on the last 10% of trials where pragmatic likelihood estimates performed poorest (trials presented in green in **A-B**). Specifically, the pragmatic likelihood estimates provide no explanatory power for these 10% of trials: Whereas speakers are biased towards one of the two candidate expressions (green dots at relative specificity 1:1 in **B**), the RSA predicts zero bias in these decisions (red dots at relative specificity 1:1 in **B**). Consistent with our hypothesis, the listener vmPFC signals are significantly correlated with the bias level in these decisions of speakers, as shown by a linear regression of the speaker's aggregate choice probabilities against the average BOLD signal extracted from the vmPFC cluster identified in Fig. 3A for each listener within these trials **(C)**. Importantly, and consistent with our whole-brain results, there is no significant difference in how well vmPFC signals can explain the speaker choices between P+ vs. P- trials, as revealed by the insignificant interaction of P+/P- × vmPFC in the regression. In stark contrast, the speaker's choice frequencies in these trials are not related to BOLD signals in other ROIs identified by either the update signal in Fig. 2C or prior probability in Fig. S9 **(D-E)**. Finally, we found that the vmPFC ROI signal performs similarly in predicting speaker behavior in these and the remaining 90% trials ($\Delta\beta = 0.008 \pm 0.013$, $P = 0.529$). Each grey and green dot represents a trial for a listener. The error bars represent the 95% confidence interval.

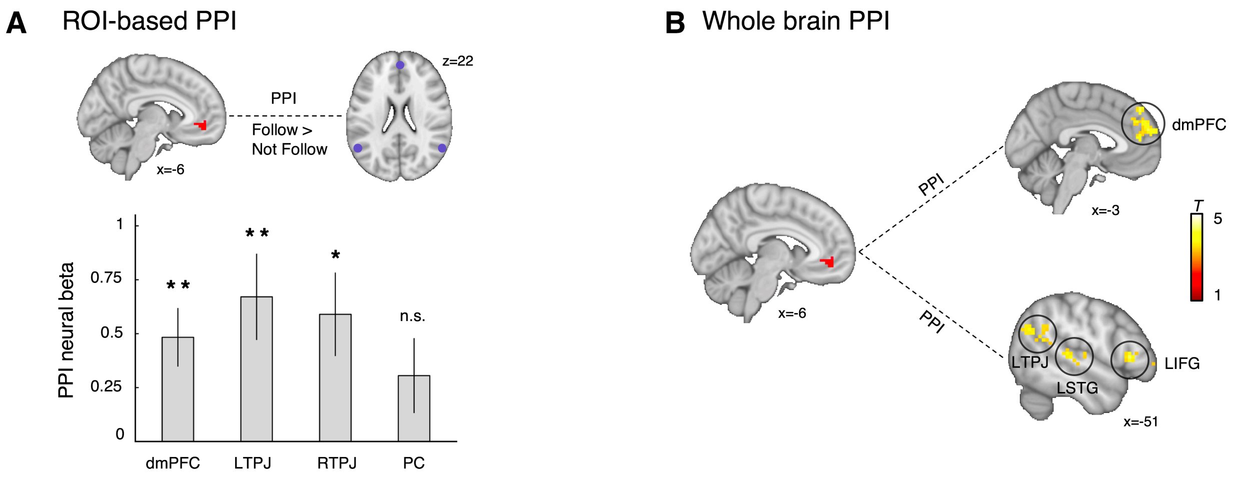


**Fig. S11.** ROI-based and explorative whole-brain PPI analyses seeded in the listener vmPFC. To explore the neural system that informs or facilitates vmPFC encoding of pragmatic likelihood estimates, we hypothesized that computing pragmatic likelihood likely requires mentalizing what the speaker would do in a given situation. Under this possibility, activity in brain regions typically implicated in mentalization, such as the dorsomedial prefrontal cortex (dmPFC) and temporoparietal junction (TPJ), may influence the encoding of pragmatic likelihood in a manner consistent with RSA predictions. To evaluate this possibility, we classified listener decisions according to whether a listener's actual choice was the one predicted by RSA with the highest posterior probability (follow model recommendations) or with a lower posterior probability (violate recommendations), and implemented a PPI analysis seeded in vmPFC during the expression onset. **(A)** Functional coupling between the vmPFC cluster identified in Fig. 3A and four theory-of-mind ROIs independently defined using Neurosyth [dmPFC, left and right TPJ, and precuneus (PC)] (*25*). In line with our prediction, there is enhanced functional coupling between vmPFC and ROIs of dmPFC and TPJ, but not PC, when the listener followed compared to when she violated RSA recommendations (dmPFC, $t_{40}$ = 3.55, $P$ = 0.004; LTPJ, $t_{40}$ = 3.35, $P$ = 0.007; RTPJ, $t_{40}$ = 3.04, $P$ = 0.016; PC, $t_{40}$ = 1.76, $P$ = 0.344; all Bonferroni corrected). **(B)** A whole-brain explorative PPI analysis with the same vmPFC cluster as the seed region also identifies the dmPFC and LTPJ, as well as three additional regions [the left inferior frontal gyrus (LIFG), the left superior temporal gyrus (LSTG), and the right dorsolateral prefrontal cortex (not shown)], demonstrate similar functional coupling patterns when listeners followed vs. did not follow model recommendations (thresholded and displayed at cluster-level $P_{\text{FWE}} < 0.05$, with a cluster-forming threshold of $P_{\text{unc.}} < 0.001$). Error bars represent inter-subject SEM. *$P < 0.05$, **$P < 0.01$, ***$P < 0.001$, all Bonferroni corrected.

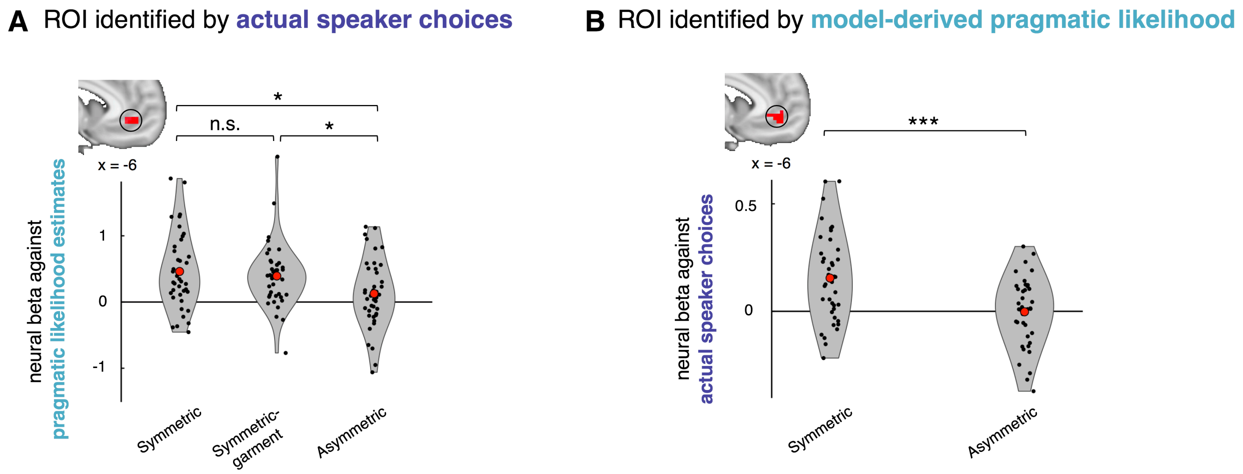


**Fig. S12.** Within-listener comparison of the neural betas in vmPFC ROIs across symmetric, symmetric-garment, and asymmetric conditions. We used two vmPFC ROIs identified by either **(A)** the actual speaker choices or **(B)** model-derived pragmatic likelihood estimates based on whole-brain search in the symmetric condition (thresholded at cluster-level $P_{\mathrm{FWE}} < 0.05$, with a cluster-forming threshold of $P_{\mathrm{unc.}} < 0.001$). For each listener in each condition, we extracted trial-wise BOLD signals from these clusters, and regressed the BOLD signal against either **(A)** the model-derived pragmatic likelihood estimates or **(B)** the speaker's actual choices. The violin plots represent the distributions of individual regression coefficients for the corresponding conditions. No correlation coefficient was computed for the symmetric-garment condition in **(B)** because no speaker data were collected for this condition (*22*). Each dot represents a listener. *$P < 0.05$, **$P < 0.01$, ***$P < 0.001$, n.s., not significant, paired-comparison, all Bonferroni corrected.

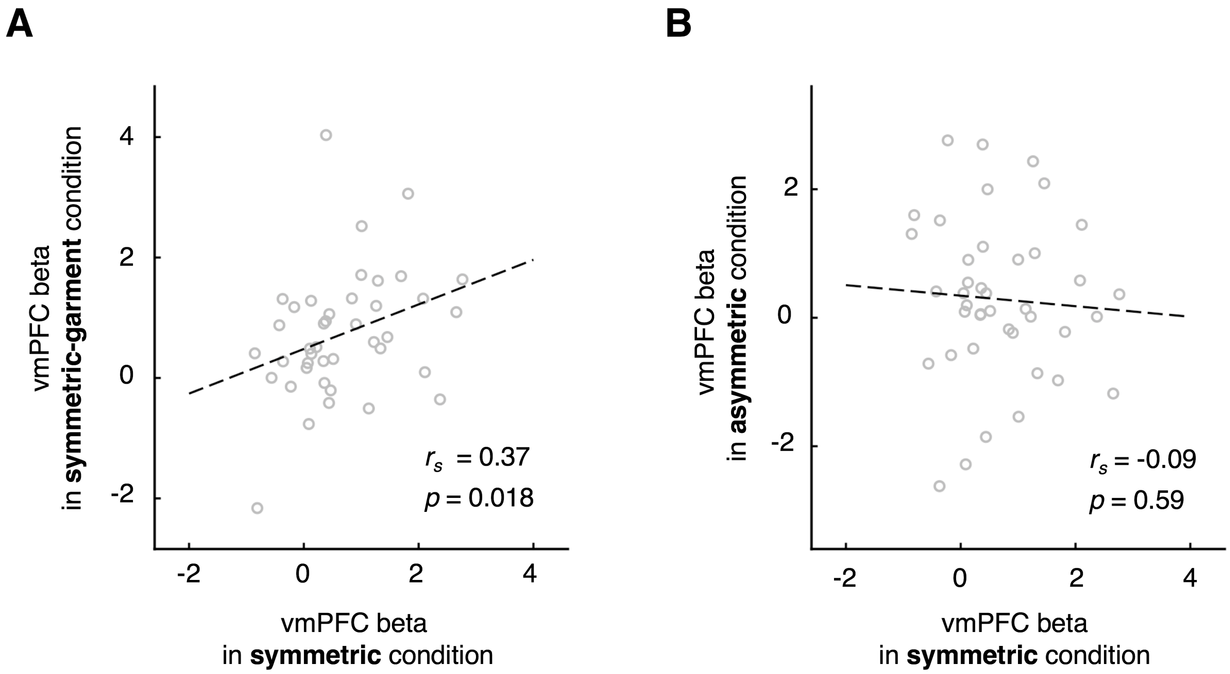


**Fig. S13.** Across-listener comparison of the neural betas extracted from the vmPFC ROI in the symmetric, symmetric-garment, and asymmetric conditions. Consistent with the finding that the vmPFC encoding of pragmatic likelihood estimates is robust to the eliciting stimuli but sensitive to the common ground information, we found positive correlation between the differential vmPFC responses to pragmatic likelihood estimates of the chosen objects in the symmetric and symmetric-garment conditions **(A)**; and no correlation between the vmPFC responses to pragmatic likelihood estimates of the chosen objects in the symmetric and asymmetric conditions **(B)**. All neural betas are extracted from the same vmPFC ROI as identified in Fig. 3A. Each dot represents a listener.

| Sign | Region | MNI peak | Voxels | T-value |
|---|---|---|---|---|
| Fig. 2C: Regions correlating with the update signal, symmetric condition | | | | |
| Positive | L/R Striatum | -9 17 -4 | 370 | 6.95 |
| | L/R Parietal/occipital lobe | 36 -67 14 | 1296 | 6.42 |
| | L/R Cerebellum | -3 -55 -22 | 347 | 6.26 |
| | L Middle temporal gyrus | -45 -76 8 | 412 | 6.17 |
| | L Thalamus | -18 -16 11 | 135 | 5.52 |
| | R Thalamus | 24 -22 23 | 61 | 5.45 |
| | R Middle frontal gyrus | 27 -4 38 | 46 | 4.98 |
| | L Caudate | -24 17 20 | 41 | 4.80 |
| | L Precentral | -33 -4 47 | 59 | 4.54 |
| Negative | R Temporal pole | 60 -22 -13 | 257 | 6.47 |
| | R Inferior parietal lobule | 54 -43 53 | 179 | 5.11 |
| | R Inferior frontal gyrus | 51 11 8 | 40 | 4.24 |
| Fig. 3A: Regions correlating with the pragmatic likelihood estimates, after controlling for trial type (P+/P-) and the posterior probability, symmetric condition | | | | |
| Positive | vmPFC | -6 44 -7 | 73 | 4.56 |
| Fig. 4B: Regions correlating with pragmatic likelihood estimates, symmetric-garment condition | | | | |
| Positive | vmPFC | 6 50 -1 | 667 | 6.03 |
| | L Temporal lobe | -33 -52 2 | 90 | 6.01 |
| | L Inferior parietal lobule | -57 -31 26 | 82 | 4.71 |
| Negative | dmPFC | -6 14 53 | 478 | 9.68 |
| | L Insula | -30 23 2 | 117 | 9.52 |
| | L/R Occipital lobe | -21 -82 -10 | 3613 | 9.16 |
| | L DLPFC | -42 11 32 | 1157 | 9.16 |
| | R DLPFC | 45 11 26 | 350 | 7.24 |
| | R Insula | 30 23 -1 | 91 | 7.02 |
| | R Superior frontal gyrus | 24 8 53 | 209 | 6.16 |
| | L Lateral PFC | -42 47 -7 | 149 | 5.99 |
| | L Caudate | -12 5 11 | 120 | 5.93 |
| | L Thalamus | -9 -16 5 | 141 | 5.91 |
| | R Thalamus | 9 -19 2 | 120 | 5.52 |
| | R Caudate | 12 5 14 | 80 | 5.22 |
| Fig. 4E: Regions correlating with pragmatic likelihood estimates derived from the matching symmetric condition, asymmetric condition. | | | | |
| Positive | R SupraMarginal | 57 -22 26 | 53 | 4.88 |

| | | | | |
|---|---|---|---|---|
| Negative | dmPFC | -6 26 38 | 586 | 8.84 |
| | L Insula | -27 23 -4 | 201 | 7.95 |
| | L DLPFC | -42 17 29 | 1154 | 7.84 |
| | L Inferior parietal lobule | -39 -49 47 | 1326 | 7.24 |
| | R Insula | 33 26 -1 | 73 | 6.89 |
| | R Superior frontal gyrus | 27 5 50 | 187 | 6.29 |
| | R DLPFC | 45 29 29 | 348 | 6.01 |
| | R Parietal lobe | 27 -61 41 | 731 | 5.62 |
| | L Caudate | -15 14 2 | 78 | 4.63 |

**Table S1.** Regions where activity correlates with trial-wise computational signals derived from the RSA model. All activations survived a cluster-level threshold $P_{\mathrm{FWE}} < 0.05$, with a cluster-forming threshold of $P_{\mathrm{unc.}} < 0.001$. L, left; R, right.
